# Correlated but distinct: VIP and NDNF interneurons implement dissociable inhibitory motifs for state-dependent gain control in visual cortex

**DOI:** 10.64898/2026.11.13.766584

**Authors:** Kyle R. Jenks, Sofie Ährlund-Richter, Marco Celotto, Grayson O. Sipe, Mriganka Sur

## Abstract

VIP and NDNF positive interneurons are both found in layers 1 and 2/3 of the cortex, but have distinct patterns of gene expression, morphology, and connectivity. Despite this, their functional roles in the sensory cortices seem remarkably similar; their activity correlates with behavioral state, and they gate behaviorally driven increases in excitatory activity and sensory-driven responses. We sought to directly compare VIP and NDNF positive interneurons in the mouse visual cortex to determine if their activity was indeed functionally redundant or could be distinguished by their correlation to behavioral state, to excitatory activity, or to excitatory visual responses. Using simultaneous recordings of pupil diameter and running speed together with dual-population calcium imaging, we found that although the activity of both VIP and NDNF positive interneurons was more strongly correlated with behavior than was excitatory neuron activity, VIP positive interneuron activity was more strongly correlated with pupil diameter, whereas layer 2/3 NDNF positive interneuron activity was more strongly correlated with running speed. Excitatory-to-VIP pairwise correlation increased during locomotion, whereas excitatory-to-NDNF pairwise correlation decreased during locomotion. Intriguingly, although VIP-to-NDNF pairwise correlation was high, cortical depth significantly affected both VIP-to-NDNF correlation and the correlation of NDNF positive interneurons with pupil diameter and running speed. As expected, excitatory visual responses were significantly modulated by both pupil dilation and locomotion. However, we found that the level of VIP and NDNF activity preceding stimulus onset had a unique contribution to and was positively correlated with the amplitude of excitatory visual responses. Thus, while highly correlated to one another and to behavioral state, VIP and NDNF positive interneurons are functionally distinguishable in their relationship to both behavior and excitatory activity in the mouse visual cortex.

## Introduction

In the visual cortex, neuronal activity, pairwise correlations, and visual responses are all modulated by behavioral state; correlating to both locomotion, as measured by running speed, and autonomic arousal, as measured by the diameter of the pupil^1–5^. Correlation of neuronal activity to behavioral state varies by cell type, with subtypes of cortical inhibitory interneurons being more correlated to behavior than excitatory neurons^3,6–9^, which provides a putative mechanism by which behavior shapes ongoing excitatory activity. There are many subtypes of cortical interneurons, and recording the activity of a specific cortical interneuron subtype of interest in vivo has largely relied on calcium imaging combined with cell-type specific molecular markers, labeling non-overlapping populations that reflect distinct morphological and electrophysiological subtypes ^8,10^. A central goal has been to attribute distinct functional roles for each inhibitory subtype in modulating excitatory activity and visual responses both during quiescence and during transitions to more active behavioral state.

Parvalbumin (PV), somatostatin (SOM), and vasoactive intestinal peptide (VIP) are non-overlapping markers of cortical inhibitory interneurons that collectively label the majority of cortical interneurons^11^. Of these subtypes, VIP positive interneurons make up the smallest proportion (roughly 15%), and are unique from PV and SOM positive interneurons in that they originate from the caudal ganglionic eminence^12^. VIP positive interneurons have a bipolar morphology with vertically elongated axons, are enriched in layer 2/3, and in visual cortex predominantly inhibit (and are inhibited by) SOM positive interneurons thereby reducing lateral inhibition^10,12–14^. This disinhibitory motif positions VIP positive interneurons as a potential pathway for state-dependent gain control, and indeed the activity of VIP positive interneurons correlates to locomotion, arousal, and both positive and negative reinforcement^7–9,13,15–17^. Moreover, ablating or chemogenetically inhibiting VIP interneurons is shown to decrease or abolish locomotion driven changes in excitatory visual responses^15,18^. Thus, one mechanism of locomotion driven gain control is hypothesized to be through excitatory or neuromodulatory input to VIP positive interneurons (but see^9^)^15^.

With improved transcriptomics able to detect increasingly sparse cell types, a fourth subtype of cortical inhibitory interneuron was identified, also originating from the caudal ganglionic eminence but largely confined to layer 1 and expressing lysosomal associated membrane protein 5 (LAMP5). While not all LAMP5 positive cells correspond to inhibitory interneurons, thus making LAMP5 ineffective without intersectional transgenic strategies, neuron-derived neurotrophic factor (NDNF) was found by multiple groups to selectively label ~70% of layer 1 inhibitory interneurons, largely overlapping with the morphologically distinct neurogliaform subtype of LAMP5 positive interneurons which are distinguished by short dendrites and a wide, dense axonal arbor^19–21^. Similar to VIP positive interneurons, NDNF positive interneurons are inhibited by SOM positive interneurons^20^. However, unlike VIP positive interneurons, NDNF positive interneurons can signal through GABAergic volume transmission, activating extrasynaptic, metabotropic GABA_B_ receptors on neighboring neurons^20,22^. In addition to inhibiting the dendrites of layer 2/3 and 5 excitatory neurons that project to L1, NDNF positive interneurons inhibit PV and VIP positive interneurons, and also presynaptically inhibit SOM positive interneurons^23–26^. While the differences in their morphology and output suggest divergent roles in regulating cortical processing, the activity of NDNF positive interneurons in vivo, and the consequences of manipulating their activity, mirror the earlier findings in VIP positive interneurons. Spontaneous activity of NDNF positive interneurons is strongly correlated with running speed and pupil diameter^27–30^. While activation of NDNF positive interneurons in anesthetized animals suppresses excitatory spontaneous activity, chemogenetic inhibition of NDNF positive interneurons in awake mice reduces the scaling of excitatory visual responses and signal-to-noise ratio with running speed and pupil diameter^27,28^. This result suggests a disinhibitory mechanism of behavioral state dependent control similar to VIP positive interneurons.

The findings from different studies suggest that VIP and NDNF positive interneurons have strong functional parallels in their relationship to behavioral state. Some studies have suggested that behavioral state modulation is largely global and non-specific, with increasing behavioral activity correlated to higher overall neuronal activity across inhibitory subtypes^6,8,9^. However, as many studies record from a single population at a time, it is unclear how the activity of VIP and NDNF positive interneurons correlates to ongoing excitatory activity and visual responses during behavior, and importantly how their activity correlates to one another. Additionally, differences in their overall correlation to pupil diameter, running speed, and excitatory activity could be lost in the variability between the approaches and criteria of the different studies. The goal of our study was to directly compare and contrast the activity of VIP and NDNF positive interneurons in the mouse visual cortex to address these gaps.

## Results

### Dual-color co-imaging of three cortical cell-type pairs in awake mice

We imaged calcium activity in the primary visual cortex of adult head-fixed mice that viewed visual stimuli on a monitor while we recorded pupil diameter and running speed (**Figure 1A**). In some sessions, we used a brief air puff to elicit shifts in behavioral state. In separate cohorts, we co-imaged three pairs of populations: VIP positive interneurons with putative excitatory neurons (VIP-Cre), NDNF positive interneurons with putative excitatory neurons (NDNF-Cre), and VIP with NDNF positive interneurons (VIP-Flp x NDNF-Cre) (**Figure 1B**). In the VIP-Cre and NDNF-Cre cohorts, pan-neuronal GCaMP7f expression was driven by the synapsin promoter while Cre-dependent tdTomato was used to label the VIP or NDNF positive populations, respectively. We refer to the unlabeled population 100 µm below the cortical surface as putative excitatory neurons but note that this may include unlabeled interneurons^31^. We refer to these cells as excitatory neurons hereafter for simplicity. We inferred firing rates from the calcium signals with Cascade and used the inferred firing rate as the measure of neuronal activity throughout^32^.

**Figure 1:**
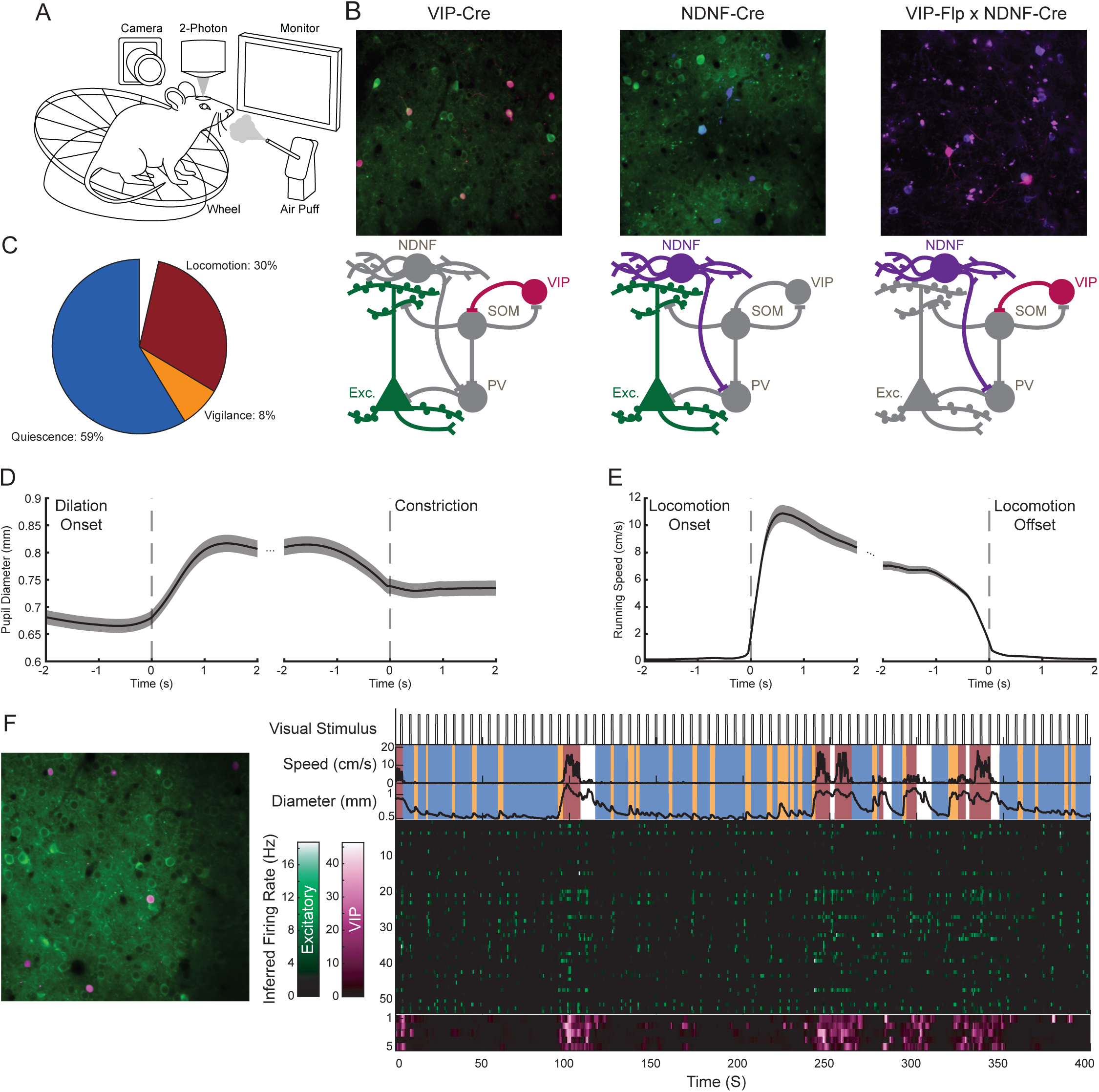
Dual-color co-imaging of three cortical cell-type pairs in awake mice. **A.** Diagram of recording setup. Head-fixed mice viewed visual stimuli on a monitor while calcium activity was imaged in the visual cortex. Pupil diameter and locomotion were recorded, and an air puff was used to elicit shifts in behavioral state. **B.** Pairs of neuronal subtypes were recorded: VIP positive interneurons and putative excitatory (excitatory) neurons (left); NDNF positive interneurons and excitatory neurons (middle); and VIP and NDNF positive interneurons (right). **C.** Behavioral state was segmented into non-overlapping states of quiescence (pupil not dilated, no locomotion), vigilance (pupil dilated, no locomotion and excluding periods following locomotion offset), and locomotion (locomotion). **D.** Labeled pupil dilation events (left) and subsequent constriction (right). **E.** Labeled locomotion event onset (left) and offset (right). N = 145 separate recordings from 13 mice. **F.** Example co-imaging of VIP interneurons and excitatory neurons. Field of view with VIP positive interneurons labeled by tdTomato (left). Section of the recording showing visual stimulus timing, running speed, pupil diameter, and a raster of excitatory (green) and VIP (magenta) inferred firing rates.

We segmented each recording into three non-overlapping behavioral states: quiescence (pupil not dilated, no locomotion), vigilance (pupil dilated, no locomotion), and locomotion (**Figure 1C**)^7^. We detected pupil dilation events and the subsequent constriction (**Figure 1D**), and locomotion onsets and offsets (**Figure 1E**), in 145 recordings from 13 mice that had both pupil and running data collected (**Figure 1E**). Across these recordings, pupil diameter and running speed covaried significantly, and changes in running speed significantly preceded changes in pupil diameter (**Figure S1A**). Pupil diameter significantly increased at locomotion onset and significantly decreased at locomotion offset (**Figure S1B**). Due to this covariance and the slower decay of pupil related activity relative to changes in locomotion, we excluded vigilance periods preceded by bouts of locomotion^33^. Pupil dilations evoked by an air puff were significantly larger than spontaneous dilations, and locomotion bouts evoked by an air puff were significantly faster than spontaneous bouts (**Figure S1C,D**). An example segment of a VIP-Cre recording shows the field of view with tdTomato-labeled VIP interneurons, together with the visual stimulus timing, running speed, pupil diameter, behavioral state, and rasters of inferred firing rates for excitatory and VIP neurons (**Figure 1F**).

### Excitatory neurons, VIP interneurons, and NDNF interneurons differ in their relationship to pupil diameter and running speed

We first asked how strongly each cell type is correlated with pupil diameter in layer 2/3. Correlation with pupil diameter differed significantly between cell types (**Figure 2A**). VIP and NDNF interneurons were each significantly more correlated with pupil diameter than excitatory neurons, and VIP interneurons were significantly more correlated with pupil diameter than NDNF interneurons (**Figure 2A**). In the cross covariance between each neuron of each cell type and pupil diameter, changes in NDNF activity significantly preceded changes in pupil diameter (**Figure 2B**). The lags between the activity of excitatory neurons and pupil diameter and between the activity of VIP interneurons and pupil diameter did not differ significantly from zero (**Figure 2B**). When we binned activity by pupil diameter decile, calculated separately for each mouse, the activity of all three cell types increased significantly with pupil diameter decile (**Figure 2C**). Next, we aligned the activity of neurons that responded to pupil dilation to dilations that occurred without locomotion. The amplitude of the normalized response differed significantly between cell types (**Figure 2D**). VIP positive interneurons had a higher pre-to-post change in amplitude than excitatory neurons, whereas the NDNF positive interneuron response did not differ significantly from the excitatory response (**Figure 2D**).

**Figure 2:**
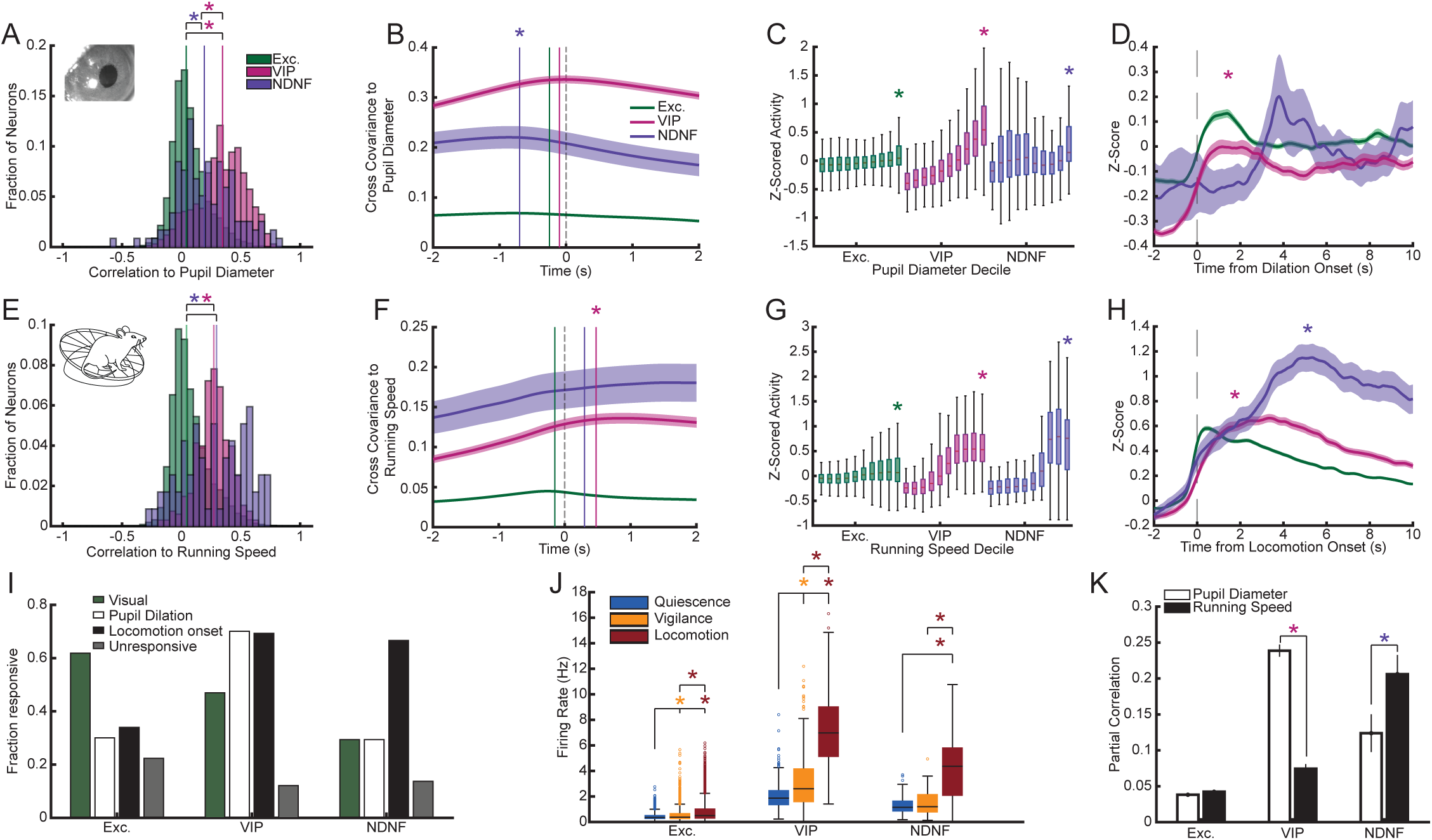
Excitatory neurons, VIP interneurons, and NDNF interneurons differ in their relationship to pupil diameter and running speed. **A.** Histogram of Pearson’s correlation of neurons to pupil diameter. Correlation to pupil is significantly different between cell types (F(2,8712.5) = 1080.8, p < 1e-300, ANOVA), with VIP and NDNF significantly more correlated to pupil than excitatory neurons (VIP: t(8840) = 45.14, p < 1e-300. NDNF: t(8840) = 11.31, p = 1.83e-29), and VIP significantly more correlated to pupil than NDNF (t(8840) = −7.550, p = 4.79e-14). **B.** Cross covariance of neurons to pupil diameter. Changes in NDNF activity precede changes in pupil diameter (z = −3.64, p = 2.76e-4, signed rank test, n = 81). **C.** Neuronal activity binned by pupil diameter decile (calculated separately for each mouse). Activity of all 3 neuron types significantly scales with pupil diameter decile (Exc.: F(1,67509) = 2333.8, p < 1e-300. VIP: F(1,5873.7) = 5631.7, p < 1e-300. NDNF: F(1,879.47) = 85.386, p = 1.8e-19, ANOVA). **D.** Averaged Z-scored activity of neurons significantly responsive to pupil dilation aligned to pupil dilations occurring in the absence of locomotion. VIP interneurons have a larger response to dilations than do excitatory neurons (Effect of cell type: F(2,1405.2) = 6.780, p = 1.17e-3, ANOVA. VIP: t(1436) = 3.500, p = 4.79e-4). **E.** Histogram of Pearson’s correlation of neurons to running speed. Correlation to running speed is significantly different between cell types (F(2,4914.4) = 292.05, p = 1.42e-120, ANOVA), with VIP and NDNF significantly more correlated to running speed than excitatory neurons (VIP: t(4980) = 21.40, p = 2.82e-97. NDNF: t(4980) = 11.32, p = 2.35e-29). **F.** Cross covariance of neurons to running speed. Changes in VIP activity lag behind changes in running speed (z = 8.21, p = 2.16e-16, signed rank test, n = 246). **G.** Activity of all 3 neuron types significantly scales with running speed decile (Exc.: F(1,41292) = 5615.5, p < 1e-300. VIP: F(1,2925) = 3673.4, p < 1e-300. NDNF: F(1,630) = 339.11, p = 6.5e-61, ANOVA). **H.** Averaged Z-scored activity of neurons responsive to locomotion onset aligned to locomotion onset. VIP and NDNF interneurons have a larger response to locomotion onset than do excitatory neurons (effect of cell type: F(2,1673.1) = 33.00, p = 8.78e-15, ANOVA. VIP: t(1738) = 7.879, p = 5.77e-15. NDNF: t(1738) = 2.009, p = 0.0447). **I.** Fraction of neurons significantly responsive to visual stimulation, pupil dilation, or locomotion (n = 3642 excitatory, 264 VIP, and 51 NDNF neurons). The fraction responsive differed by cell type for visual stimulation (F(2,3954) = 24.02, p = 4.28e-11), pupil dilation (F(2,3954) = 67.19, p = 2.02e-29), and locomotion (F(2,3954) = 70.63, p = 7.29e-31; binomial generalized linear mixed-effects model). Compared with excitatory neurons, fewer VIP (t(3954) = −6.147, p = 8.67e-10) and NDNF (t(3954) = −3.218, p = 1.30e-3) neurons responded to visual stimulation, more VIP neurons responded to pupil dilation (t(3954) = 11.57, p = 1.79e-30), and more VIP (t(3954) = 11.04, p = 6.00e-28) and NDNF (t(3954) = 4.449, p = 8.86e-6) neurons responded to locomotion. NDNF and excitatory neurons did not differ in the fraction responsive to pupil dilation (t(3954) = 0.773, p = 0.439). **J.** Average neuronal firing rate during each behavioral state. Excitatory firing rate significantly changes between behavioral states (F(2,9176) = 777.69, p < 1e-300, ANOVA), increasing from quiescence to vigilance (t(13761) = 11.82, p = 4.55e-32) and vigilance to locomotion (t(13761) = 26.68, p = 6.56e-153). VIP firing rate significantly changes between behavioral states (F(2,650) = 776.6, p = 5.06e-173, ANOVA), increasing from quiescence to vigilance (t(972) = 7.623, p = 5.87e-14) and vigilance to locomotion (t(972) = 29.67, p = 2.73e-138). NDNF firing rate significantly changes between behavioral states (F(2,210) = 75.442, p = 2.04e-25, ANOVA), increasing from quiescence to locomotion (t(207) = 10.888, p = 4.03e-22) and from vigilance to locomotion (t(207) = 10.368, p = 1.48e-20), but not from quiescence to vigilance. **K.** Partial Correlation of neurons to pupil diameter and running speed. VIP interneurons have significantly higher partial correlation to pupil diameter than to running speed (F(1,585.09) = 257.49, p = 2.74e-48, ANOVA, n = 325 neurons), while NDNF interneurons have significantly higher partial correlation to running speed than to pupil diameter (F(1,125.93) = 5.21, p = 0.024, ANOVA, n = 70 neurons). *denotes significance.

We next examined each cell type’s correlation to running speed. Correlation with running speed also differed significantly between cell types (**Figure 2E**). VIP and NDNF interneurons were each significantly more correlated with running speed than excitatory neurons, but, unlike pupil diameter, VIP and NDNF positive interneurons did not differ significantly from each other (**Figure 2E**). In the cross covariance with running speed, changes in VIP positive interneuron activity significantly lagged behind changes in running speed (**Figure 2F**). The lags of excitatory neurons and NDNF positive interneurons did not differ significantly from zero (**Figure 2F**). The activity of all three cell types increased significantly across running-speed deciles (**Figure 2G**). For neurons that responded to locomotion onset, the normalized response aligned to onset showed a significant effect of cell type (**Figure 2H**). VIP and NDNF positive interneurons each responded significantly more strongly than excitatory neurons, but VIP and NDNF positive interneurons did not differ significantly from each other (**Figure 2H**).

An unexpected air puff evokes a strong behavioral and neuronal response^13,17^. The amplitude of the normalized puff response differed significantly between cell types: VIP positive interneurons responded significantly more strongly than excitatory neurons, but NDNF positive interneurons did not differ significantly from excitatory neurons (**Figure S2A**). The pupil response to the puff changed significantly over the course of the first session with pupil responses significantly decreasing with repeated puffs, while running speed following puff significantly increased. The responses of excitatory neurons, VIP positive interneurons, and NDNF positive interneurons to repeated puffs each decreased significantly over the length of a recording (**Figure S2D**).

We then asked what fraction of each population was significantly responsive to visual stimulation, pupil dilation, and/or locomotion onset. A larger fraction of excitatory neurons was visually responsive than of VIP or NDNF positive interneurons. A larger fraction of VIP positive interneurons was responsive to pupil dilation than of excitatory neurons or NDNF positive interneurons, whereas the fractions of NDNF positive interneurons and excitatory neurons were not significantly different. A larger fraction of both VIP and NDNF positive interneurons was responsive to locomotion onset than of excitatory neurons (**Figure 2I**). Firing rates of all three cell types differed between behavioral states (**Figure 2J**): excitatory and VIP firing rates increased from quiescence to vigilance and further during locomotion, whereas NDNF firing rates increased only during locomotion.

Because pupil diameter and running speed covary (**Figure S1A**), correlations with either measure partly reflect the other. To ask which behavioral measure each population preferentially tracks, we computed partial correlations of each cell type with pupil diameter and with running speed. VIP positive interneurons had a significantly higher partial correlation with pupil diameter than with running speed, whereas NDNF positive interneurons had a higher partial correlation with running speed than with pupil diameter (**Figure 2K**). Excitatory neurons did not differ significantly in their partial correlation with the behavioral measurements (**Figure 2K**). Thus, our results suggest that while both VIP and NDNF positive interneurons are more correlated and responsive to behavioral state and behavioral state transitions than are excitatory neurons, VIP activity preferentially tracks pupil diameter, whereas NDNF activity preferentially tracks running speed.

### Coupling between excitatory, VIP, and NDNF populations depends on behavioral state and distance

Having found that VIP and NDNF positive interneurons differ in how they track behavior, we next asked how their mean activity covaries with both mean excitatory activity and with one another. In VIP-Cre recordings, the peak of the cross covariance lagged such that changes in mean excitatory activity significantly preceded changes in mean VIP activity (**Figure 3A**). In NDNF-Cre recordings, the lag did not differ significantly from zero. In VIP-Flp x NDNF-Cre recordings, changes in mean NDNF activity significantly preceded changes in mean VIP activity (**Figure 3A**). When we separated the cross covariance by behavioral state, the peak covariance between mean excitatory activity and mean NDNF activity differed significantly between states, decreasing during periods of locomotion (**Figure 3C**). We then asked how pairwise correlation of individual neurons with neurons of the other population depends on state. The mean pairwise correlation of each excitatory neuron with the co-imaged VIP positive interneurons differed significantly between behavioral states and was significantly higher in locomotion than in quiescence but did not differ significantly between vigilance and quiescence (**Figure 3E, top**). The mean pairwise correlation of each excitatory neuron with the co-imaged NDNF positive interneurons also differed significantly between states: it was significantly higher in vigilance than in quiescence but significantly lower in locomotion than in quiescence (**Figure 3E, middle**). The mean pairwise correlation of each VIP positive interneuron with the co-imaged NDNF positive interneurons differed significantly between states and was significantly higher in vigilance and in locomotion than in quiescence (**Figure 3E, bottom**). We noted that in the VIP-Flp x NDNF-Cre data, where imaging was confined to layer 1, the correlation between VIP positive interneurons and NDNF positive interneurons shallower than 50 µm was significantly lower than the correlation between VIP positive interneurons and NDNF positive interneurons at 50 µm or deeper (**Figure 3F**).

**Figure 3:**
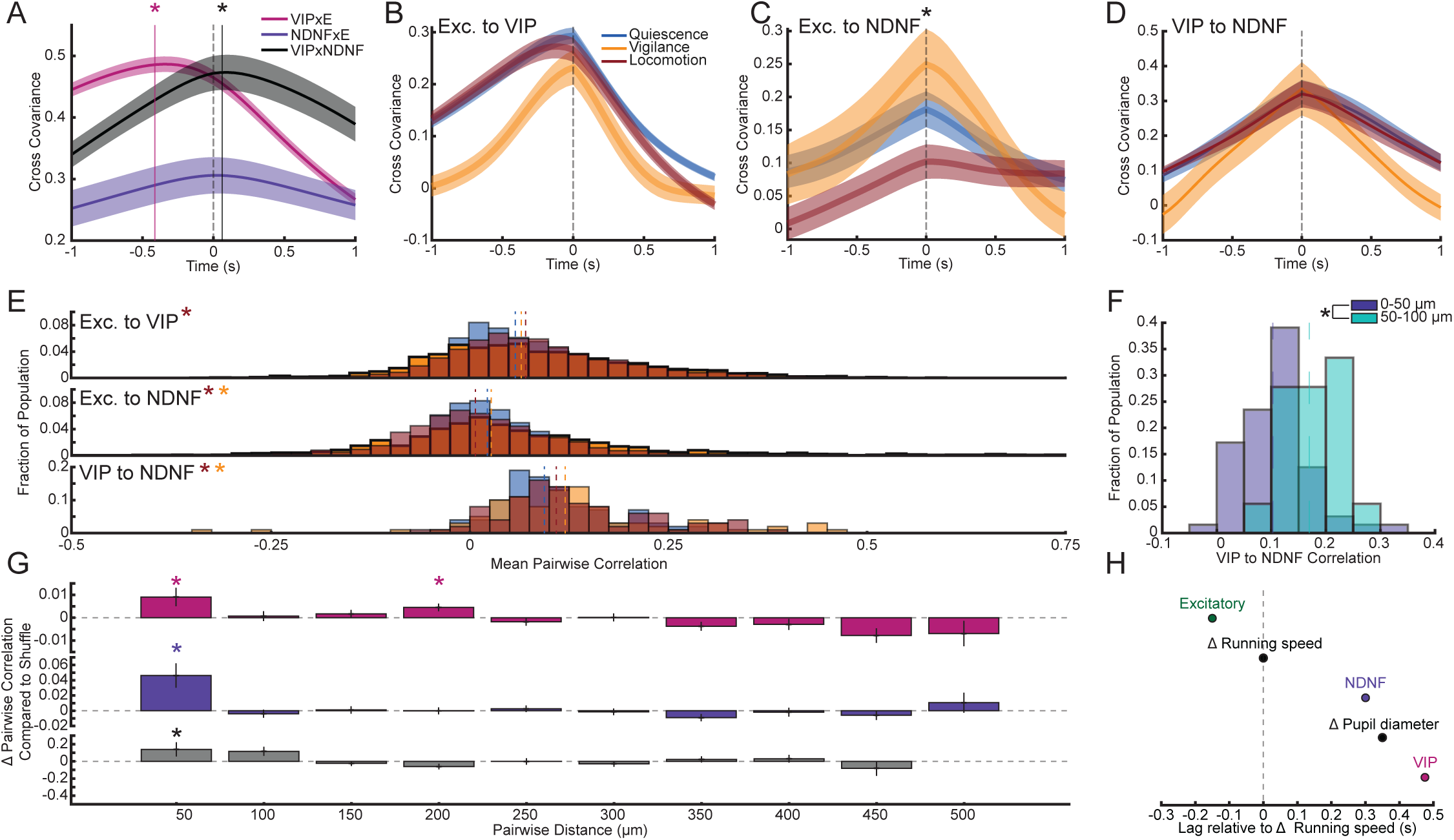
Coupling between excitatory, VIP, and NDNF populations depends on behavioral state and distance. **A.** Cross covariance of co-imaged populations. Mean excitatory neuron and VIP interneuron activity significantly covary (z(118) = 9.427, p = 4.21e-21, signed rank test), and changes in excitatory activity precede changes in VIP activity (z(118) = −9.062, p = 1.28e-19, signed rank test). Mean excitatory neuron and NDNF interneuron activity significantly covary (z(31) = 4.86, p = 1.17e-6, signed rank test). Mean VIP and NDNF interneuron activity significantly covary (z(17) = 3.621, p = 2.93e-4, signed rank test), and changes in NDNF activity precede changes in VIP activity (z(17) = 2.132, p = 0.033, signed rank test). **B.** Cross covariance of mean excitatory neuron and VIP interneuron activity separated by behavioral state. **C.** Cross covariance of mean excitatory neuron and NDNF interneuron activity separated by behavioral state. Excitatory to NDNF peak covariance significantly changes between behavioral states (F(2,35.159) = 7.43, p = 2.04e-3, ANOVA). **D.** Cross covariance of VIP and NDNF interneurons separated by behavioral state. **E.** (Top) Histogram of each excitatory neuron’s mean correlation to all co-imaged VIP interneurons. Correlation significantly changes from quiescence to locomotion (F(2,6508) = 4.745, p = 8.72e-3, ANOVA for effect of behavioral state. Locomotion vs. quiescence: t(9759) = 3.080, p = 2.07e-3). (middle) Histogram of each excitatory neuron’s mean correlation to all co-imaged NDNF interneurons. Correlation significantly changes between behavioral states (F(2,2668) = 37.349, p = 1.01e-16, ANOVA), increasing from quiescence to vigilance (t(3999) = 3.221, p = 1.29e-3) and decreasing from quiescence to locomotion (t(3999) = −5.336, p = 1e-7). (Bottom) Histogram of each VIP interneuron’s mean correlation to all co-imaged NDNF interneurons. Correlation significantly changes between behavioral states (F(2,200) = 3.112, p = 0.0467, ANOVA), increasing from quiescence to vigilance (t(297) = 2.124, p = 0.0345) and quiescence to locomotion (t(297) = 2.195, p = 0.029). **F.** VIP interneurons’ mean correlation to co-imaged NDNF interneurons is significantly different within 0-50 and 50-100 µm from the cortical surface (F(1,13.19) = 13.206, p = 2.96e-3, ANOVA). **G.** Histograms of pairwise correlations between co-imaged neurons binned by their horizontal distance apart and plotted as the delta correlation of the observed mean correlation compared to the mean correlation of the shuffled data. (Top) Excitatory-to-VIP correlation was significantly higher than the shuffled distribution between 0-50 (p = 0.026, permutation test, 10,000 shuffles, Bonferroni-corrected, n=1176 pairs, N=118 FOVs) and 150-200 µm apart (p = 6e-3, permutation test, 10,000 shuffles, Bonferroni-corrected, n=5361 pairs, N=119 FOVs). (Middle) Excitatory-to-NDNF correlation was significantly higher than expected at 0-50 µm apart (p < 2e-3, permutation test, 10,000 shuffles, Bonferroni-corrected, n=172 pairs, N=32 FOVs). (Bottom) VIP-to-NDNF correlation was significantly higher than expected from 0-50 µm apart (p = 4e-3, permutation test, 10,000 shuffles, Bonferroni-corrected, n=23 pairs, N=7 FOVs). **H.** Diagram representing the order in which signals change, relative to the lag in their cross covariance to running speed. Note that the change in running speed is being used as a common reference point for the other signals, and as such this does not represent the true lag in those signals’ cross covariance to one another, which is dependent on factors other than changes in running speed. *denotes significance.

This result raised the question of whether NDNF positive interneuron activity depends on depth. Indeed, NDNF positive interneuron subtype varies with cortical depth, and LAMP5 positive interneurons differ in their connectivity and correlation to locomotion between layers 1 and 2/3^21,28^. The NDNF-Cre recordings analyzed so far were restricted to recordings deeper than 100 µm, to better match the excitatory and VIP positive interneuron data, and so we repeated the analyses of firing rate, correlation, and partial correlation in all NDNF-Cre recordings, including those shallower than 100 µm, divided into three depth categories (0 to 50 µm, 50 to 100 µm, and more than 100 µm; **Figure S3**). The dependence of NDNF firing rate on behavioral state differed significantly between depth categories (**Figure S3A**). At 0 to 50 µm, NDNF firing rates in vigilance and in locomotion were each significantly higher than in quiescence, and firing in locomotion was significantly higher than in vigilance (**Figure S3A**). At 50 to 100 µm and at more than 100 µm, firing rate increased significantly in locomotion relative to quiescence and relative to vigilance but did not differ significantly between quiescence and vigilance (**Figure S3A**). The correlation of NDNF positive interneurons with pupil diameter differed significantly between depth categories and was significantly higher at depths greater than 100 µm than at 0 to 50 µm or 50 to 100 µm (**Figure S3B**). The correlation with running speed also differed significantly between depth categories and was significantly higher at depths greater than 100 µm than at 0 to 50 µm or 50 to 100 µm (**Figure S3C**). In the partial correlations, NDNF positive interneurons differed significantly between pupil diameter and running speed in each of the three depth categories. Partial correlation with running speed was greater than partial correlation with pupil diameter at 0 to 50 µm and at more than 100 µm; the reverse was true at 50 to 100 µm (**Figure S3D**). NDNF positive interneurons at 50 to 100 µm appear to be a functionally distinct subset, less driven by changes in locomotion and more driven by changes in pupil, corroborating the higher VIP-to-NDNF correlation observed at 50 to 100 µm.

We also asked whether interpopulation correlation depends on the lateral distance between neurons, which has been demonstrated previously for spontaneous excitatory-to-VIP correlations^14^. We binned pairwise correlations by the distance between the two neurons and compared the observed mean correlation in each bin with a shuffled distribution, with correction for the number of bins (**Figure 3G**). Excitatory-to-VIP correlation was significantly higher than the shuffled distribution at distances of 0 to 50 µm and 150 to 200 µm, and not significantly different at the other distances (**Figure 3G, top**). Excitatory-to-NDNF correlation was significantly higher than the shuffled distribution at 0 to 50 µm only (**Figure 3G, middle**). VIP-to-NDNF correlation was significantly higher than the shuffled distribution at 0 to 50 µm only (**Figure 3G, bottom**). The overall lag in changes of mean population activity to one another is likely driven in part by their varying lags to changes in behavioral state, which we visualize here by aligning the peak of cross covariance of excitatory, VIP interneuron, NDNF interneuron, and pupil activity to changes in running speed (**Figure 3H**). Because changes in running speed serve as a common reference, the diagram does not accurately reflect the lag between the populations themselves, the variance of which is driven by factors other than locomotion. In summary, we found that both VIP and NDNF positive interneurons are positively correlated with excitatory neurons, but locomotion changes these correlations in opposite directions, increasing them for VIP and decreasing them for NDNF positive interneurons. We also found that NDNF positive interneurons’ correlation to both neighboring VIP positive neurons and behavioral measures varies by depth.

### Excitatory visual responses scale with contrast and behavioral state

Locomotion and arousal have been shown to uniquely contribute to changes in excitatory visual responses^3,7^. Therefore, we sought to quantify how this behavioral modulation was reflected in our segmentation of behavioral state. As we were interested in behavioral state as a modulator of visual response, as opposed to any possible behavioral change driven by the stimulus itself, we assigned a given trial’s behavioral state as the behavioral state recorded at the onset of the stimulus and measured visual response as the change in firing rate from the pre-stimulus period, to account for behaviorally driven changes in mean firing rate (**Figure 2J**). Responses of excitatory neurons to drifting gratings, aligned to each neuron’s preferred direction at 64% contrast, scaled significantly with contrast (**Figure 4A**). Behavioral state also had a significant effect, and the interaction between contrast and behavioral state was significant. At 4% contrast, responses during locomotion were significantly higher than during quiescence, and responses during vigilance did not differ significantly from quiescence (**Figure 4B**). At 16% contrast, responses during vigilance and during locomotion were each significantly higher than during quiescence (**Figure 4C**). At 64% contrast, responses during locomotion were significantly higher than during quiescence, and responses during vigilance did not differ significantly from quiescence (**Figure 4D**). We next tested whether the effects of behavior were greater for specific directions in the visual field. We plotted normalized responses to 64% contrast gratings by the direction in which the grating moved on the monitor, separated by behavioral state (**Figure 4E**). Direction had a significant effect overall, and responses to 45° and 135° were significantly higher than responses to 0° (**Figure 4E**). The interaction between direction and behavioral state was significant (**Figure 4E**). These two directions align with the axis of expected visual flow, with 45° moving opposite to the expected visual flow for forward running.

**Figure 4:**
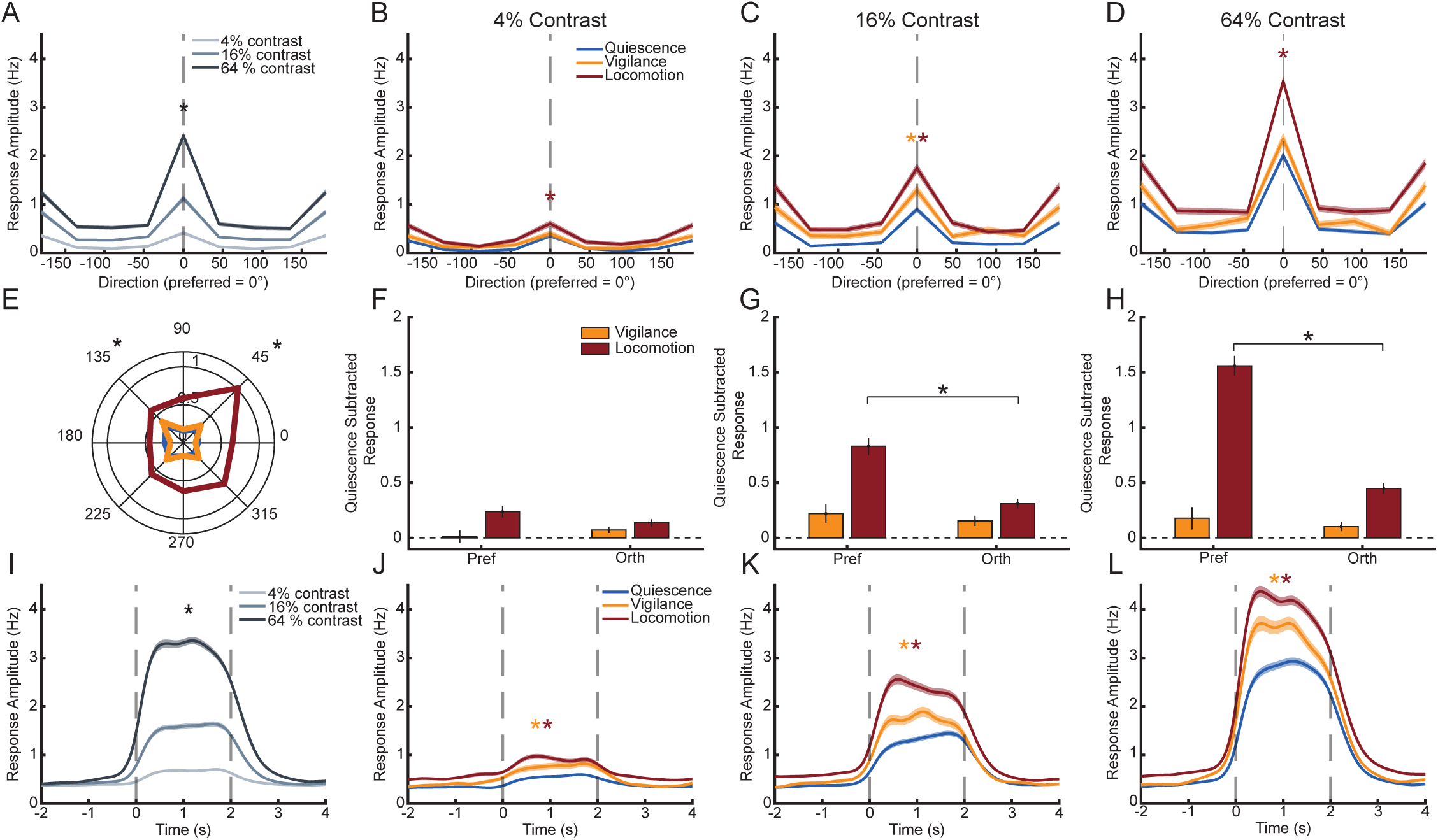
Excitatory visual responses scale with behavioral state. **A.** Excitatory responses to drifting gratings scale with contrast (F(2,5379) = 14.864, p = 3.65e-7, ANOVA). Responses of each neuron are aligned to their preferred direction at 64% contrast. There is an additional significant effect of behavior (F(2,5466.4) = 3.158, p = 0.043, ANOVA) and contrast:behavioral state interaction (F(4,5443.8) = 5.476, p = 2.13e-4, ANOVA). **B.** Responses at 4% contrast to drifting gratings separated by behavioral state at the onset of the visual stimulus. Responses during locomotion are higher than responses during quiescence (t(2147) = 4.75, p = 2.14e-6). **C.** Same as B., but for 16% contrast. Responses during vigilance (t(2045) = 2.593, p = 9.57e-3) and locomotion (t(2045) = 10.598, p = 1.39e-25) are higher than responses during quiescence. **D.** Same as B., but for 64% contrast. Responses during locomotion are higher than responses during quiescence (t(2062) = 17.666, p = 3.79e-65). **E.** Normalized excitatory visual responses to 64% contrast gratings aligned to the direction the grating was moving on the monitor, separated by behavioral state. There was a significant effect of direction overall (F(7,2569) = 5.454, p = 3.12e-6, ANOVA), with 45° (t(2928) = 2.39, p = 0.0169) and 135° (t(2928) = 3.468, p = 5.33e-4) significantly different than 0°. There was a significant interaction between direction and behavior (F(14,6587.3) = 4.825, p = 6.01e-9). **F.** Each neuron’s responses to gratings during quiescence were subtracted from its responses to the same stimulus during vigilance and locomotion states, and the interaction between behavioral state and direction type (preferred (pref) vs orthogonal (orth)) was tested. There was a significant interaction between preferred direction, contrast, and behavioral state (F(4,12420) = 11.894, p = 1.21e-9, ANOVA). At 4% contrast, the interactions between behavioral state and preferred direction were not significant. **G.** Same as F., but for 16% contrast. The interaction between the locomotion state and preferred direction was significant (t(4397) = 5.443, p = 5.54e-8). **H.** Same as F., but for 64% contrast. The interaction between the locomotion state and preferred direction was significant (t(4352) = 10.103, p = 9.815e-24). **I.** Excitatory responses to natural movies scale with contrast (F(2,8494.7) = 143.18, p = 6.93e-62, ANOVA). Responses of each neuron are to their preferred movie at 64% contrast. There is an additional significant contrast:behavioral state interaction (F(4,8595.6) = 10.281, p = 2.65e-8, ANOVA). **J.** Responses at 4% contrast to natural movies separated by behavioral state at the onset of the visual stimulus. Responses during vigilance (t(3196) = 3.477, p = 5.15e-4) and locomotion (t(3196) = 3.853, p = 1.19e-4) are higher than responses during quiescence. **K.** Same as J., but for 16% contrast. Responses during vigilance (t(3294) = 3.162, p = 1.58e-3) and locomotion (t(3294) = 15.06, p = 1.29e-49) are higher than responses during quiescence. **L.** Same as J., but for 64% contrast. Responses during vigilance (t(3411) = 6.556, p = 6.36e-11) and locomotion (t(3411) = 17.893, p = 1.64e-68) are higher than responses during quiescence. *denotes significance.

To test whether state effects are specific to the preferred direction, we subtracted each neuron’s mean response in quiescence from its mean responses in vigilance and locomotion at both the neuron’s preferred and orthogonal directions. The three-way interaction among preferred versus orthogonal direction, contrast, and behavioral state was significant (**Figure 4F**). At 4% contrast, the interactions between behavioral state and direction type were not significant (**Figure 4F**). At 16% and 64% contrast, the interaction between locomotion and direction type was significant, with a larger locomotion-related increase for the preferred direction (**Figure 4G,H**). The interaction between vigilance and direction type was not significant at any contrast. Thus, visual responses were highest during locomotion across all contrasts, and locomotion sharpened selectivity through increasing preferred responses more than orthogonal responses, consistent with multiplicative gain. Responses to natural movies, measured for each neuron’s preferred movie at 64% contrast, also scaled significantly with contrast, and the interaction between contrast and behavioral state was significant (**Figure 4I**). At each of the three contrasts, responses during vigilance and during locomotion were significantly higher than during quiescence (**Figure 4J–L**). Thus, while visual responses to gratings and natural movies are positively modulated by locomotion, modulation by vigilance is most evident for responses to more naturalistic stimuli.

### Excitatory visual responses scale with VIP and NDNF activity beyond behavioral state measures

We next asked whether the activity of VIP and NDNF positive interneurons at stimulus onset relates to excitatory visual responses, beyond their correlation with behavioral state. For each stimulus trial in quiescence, we classified the preceding interneuron state by the quartile of mean (non-zero) VIP positive interneuron activity at stimulus onset (**Figure 5A**) and again took the amplitude of the visual response as the change in firing rate from pre- to post-stimulus to account for elevation of mean excitatory activity correlated with VIP activity (**Figure 3A**). Excitatory responses to each neuron’s preferred grating scaled significantly with contrast, and the interaction between contrast and VIP activity quartile was significant (**Figure 5B**). Response amplitude varied significantly with VIP activity quartile at each of the three contrasts (**Figure 5B**). For natural movies (**Figure 5C**), as with gratings, the interaction between contrast and VIP activity quartile was significant (**Figure 5D**). Response amplitude varied significantly with VIP activity quartile at 16% and 64% contrast, and not significantly at 4% contrast (**Figure 5D**). We repeated the analysis with quartiles of mean NDNF positive interneuron activity (**Figure 5E**). For gratings, the interaction between contrast and NDNF activity quartile was significant, and response amplitude varied significantly with NDNF activity quartile at 4% and 64% contrast, but not at 16% contrast (**Figure 5F**). For natural movies (**Figure 5G**), the interaction between contrast and NDNF activity quartile did not reach significance (**Figure 5H**).

**Figure 5:**
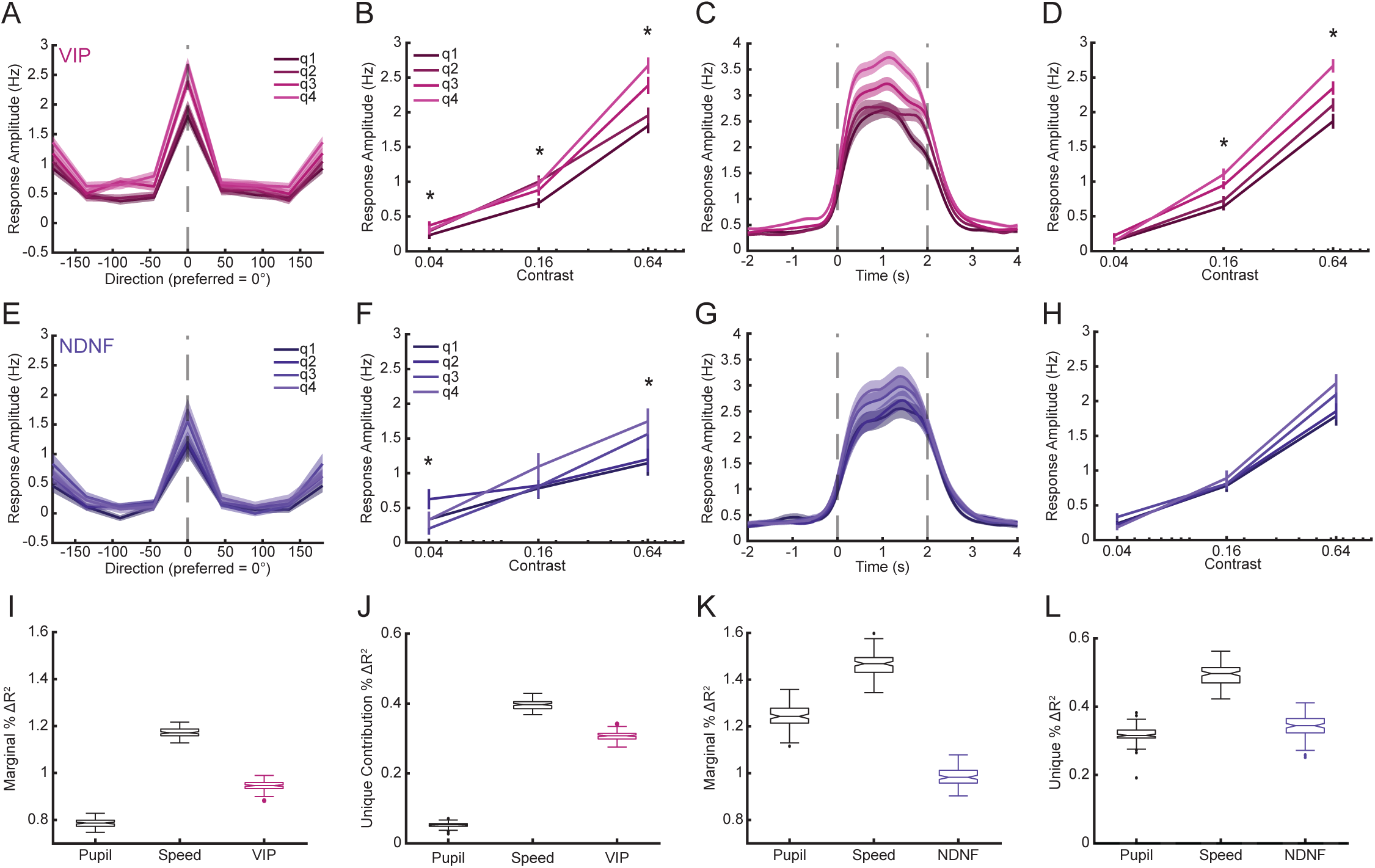
Excitatory visual responses scale with VIP and NDNF activity beyond behavioral state measures. **A.** Response amplitude (post-pre) of excitatory neurons to 64% contrast drifting gratings during quiescence, with each neuron’s responses aligned to their preferred direction. For each stimulus trial, the quartile of mean VIP interneuron activity at the onset of the stimulus is used to categorize the background interneuron state during which the trial occurred. **B.** Response amplitude of neurons to their preferred direction during quiescence scales with contrast, and there is a significant interaction of contrast and the quartile of mean VIP interneuron activity at stimulus onset (F(6,5804.8) = 7.707, p = 2.86e-8, ANOVA). Response scales with VIP activity quartile at all contrasts (4%: F(3,1470.1) = 2.67, p = 0.0462. 16%: F(3,1514.7) = 3.43, p = 0.0165. 64%: F(3,1522.6) = 20.009, p = 1.013e-12. ANOVA). **C.** Response amplitude of excitatory neurons to 64% contrast natural movies during quiescence, with each neuron’s responses aligned to their preferred natural movie. For each stimulus trial, the quartile of mean VIP interneuron activity at the onset of the stimulus is used to categorize the background interneuron state during which the trial occurred. **D.** Response amplitude of neurons to their preferred natural movie during quiescence scales with contrast, and there is a significant interaction of contrast and the quartile of mean VIP interneuron activity at stimulus onset (F(6,7687.9) = 8.425, p = 3.91e-9, ANOVA). Responses scale with VIP quartile at 16% (F(3,2001.9) = 12.367, p = 5.14e-8, ANOVA) and 64% (F(3,1920.9) = 38.794, p = 2.45e-24, ANOVA) contrast. **E.** Same as A., but binning by mean NDNF interneuron activity quartile. **F.** Response amplitude of neurons to their preferred direction during quiescence scales with contrast, and there is a significant interaction of contrast and the quartile of mean NDNF interneuron activity at stimulus onset (F(6,1400.3) = 2.447, p = 0.0234, ANOVA). Responses scale with NDNF activity quartile at 4% (F(3,383.19) = 3.387, p = 0.0182, ANOVA) and 64% contrast (F(3,367.97) = 4.61, p = 3.51e-3, ANOVA). **G.** Same as C., but binning by mean NDNF interneuron activity quartile. **H.** Response amplitude of neurons to their preferred natural movie during quiescence scales with contrast, but the interaction between contrast and NDNF activity quartile does not reach significance (F(6,3330.3) = 2.0314, p = 0.0582, ANOVA). **I.** Mixed-effect linear models were fit to the amplitude of excitatory visual responses with contrast as a predictor, and either the pupil diameter, running speed, or mean VIP interneuron activity preceding stimulus onset (and their interaction with contrast) as predictors (52,560 trials, 1,752 neurons, 59 sessions, 5 mice). Across 100 rounds of three-fold cross validation, the R2 of the model with just contrast was subtracted from the R2 of each of the other models. The significance of the marginal contribution of each predictor was computed in comparison to the model with just contrast using the full dataset. The marginal contributions of pupil diameter, running speed, and mean VIP activity were all significant (Pupil Diameter: p = 3.5e-148. Running Speed: p = 9.2e-223. VIP Activity: p = 3.0e-180, likelihood ratio test). **J.** The same 100 rounds of three-fold cross validation were run, but this time comparing the total explained variance of the full model with all predictors to the explained variance of models with each of the predictors removed, to estimate the unique fraction of total explained variance contributed by each predictor. The significance of the unique explained variance of each predictor compared to the full model was computed using the full dataset. Pupil diameter, running speed, and mean VIP activity preceding stimulus onset all have significant unique contributions to the explained variance (Pupil Diameter: p = 5.27e-12. Running Speed: p = 4.4e-78. VIP Activity: p = 5.5e-61, likelihood ratio test). **K.** Same as I., but for mean NDNF activity preceding stimulus onset (16,350 trials, 545 neurons, 20 sessions, 4 mice). The marginal contributions of pupil diameter, running speed, and mean NDNF activity were all significant (Pupil Diameter: p = 4.3e-65. Running Speed: p = 1.2e-82. NDNF Activity: p = 6.2e-56, likelihood ratio test). **L.** Same as J., but for mean NDNF activity preceding stimulus onset. Pupil diameter, running speed, and mean NDNF activity preceding stimulus onset all have significant unique contributions to the explained variance (Pupil Diameter: p = 7.8e-17. Running Speed: p = 5.3e-31. NDNF Activity: p = 5.5e-22, likelihood ratio test). *denotes significance.

Even during quiescence, the scaling of excitatory responses with preceding VIP and NDNF activity may reflect shared correlation with subthreshold changes in pupil diameter or locomotion. To determine if VIP and NDNF activity contributes to excitatory visual responses beyond their correlation to behavioral state, we fit mixed-effects models to the amplitude of excitatory visual responses, with contrast as a predictor and either pupil diameter, running speed, or mean interneuron activity before stimulus onset (and its interaction with contrast) as an additional predictor. In VIP-Cre recordings, each of the three predictors (pupil diameter, running speed, and mean VIP interneuron activity) made a significant marginal contribution relative to the model with contrast alone (**Figure 5I**). We then compared a model that contained all predictors with models from which one predictor was removed. We first assessed explained variance of the full model with 100 repartitions of three-fold cross-validation, and the full model explained held-out variance in both the VIP-Cre and NDNF-Cre cohorts (**Figure S4**). Pupil diameter, running speed, and VIP activity each made a significant unique contribution to the explained variance (**Figure 5J**). In NDNF-Cre recordings, pupil diameter, running speed, and NDNF activity each made a significant marginal contribution (**Figure 5K**) and a significant unique contribution (**Figure 5L**). We conclude that both VIP and NDNF activity preceding stimulus onset contributes positively to the amplitude of excitatory visual responses, beyond what can be accounted for by their correlation to behavioral state.

## Discussion

In this study, we compared and contrasted the activity of two well-studied subtypes of cortical inhibitory interneurons in the mouse visual cortex: VIP and NDNF positive interneurons. Specifically, we have compared their activity as it relates to behavioral state, excitatory activity, and excitatory visual responses. Our results concur with previously published data reporting that both subtypes are more correlated to behavioral state than excitatory neurons, but demonstrate that they are distinguished by their differential correlation to pupil diameter and running speed, two closely related but distinct measures of behavioral arousal^3,7^. While behavioral state is a powerful predictor of both excitatory activity and visual responses, both mean VIP and NDNF activity preceding stimulus onset contribute unique predictive power beyond their correlation to pupil diameter and running speed to shape excitatory responses. Thus, VIP and NDNF interneurons are not functionally redundant: they differ in their relative tracking of pupil diameter and running speed and in how their correlation with excitatory neurons changes with locomotion, while both predict larger excitatory visual responses beyond these behavioral state markers.

Although previously attempted, state-dependent modulation of excitatory visual responses has not fit a simple labeled line model driven by a singular input or cell type. This complexity is likely due to a confluence of global and local factors. The signals driving state-dependent transitions in the visual cortex include diffuse neuromodulatory inputs, such as acetylcholine and noradrenaline^6,15,33^. Locomotion leads to feedback from primary and secondary motor regions, which are important for visuomotor integration^34,35^. Behavioral state also alters activity upstream of the visual cortex in the dorsal lateral geniculate nucleus of the thalamus, and downstream of the visual cortex in regions that provide feedback^2,36^. The presence and properties of visual stimuli also dramatically alter the cell-type specific effects of behavioral state^8,9,15^. State-dependent modulation, therefore, is occurring network wide in a highly recurrent system. Reducing this complexity to a single cell-type that translates behavioral state into changes in excitatory activity is an attractively simple conceptual framework. VIP positive interneurons have been a candidate to serve this role. Optogenetic activation of VIP positive interneurons mimicked the effects of locomotion on excitatory activity, while photolytic damage, chemogenetic suppression, or developmental perturbations of VIP positive interneurons decreased state-dependent modulation of excitatory/non-VIP positive neurons^15,18,37,38^. However, other studies ablating or silencing VIP positive interneurons have shown no effect on locomotion induced changes in excitatory activity^28,39^.

NDNF positive interneurons have been another candidate cell type to link changes in behavior to changes in excitatory activity. Optogenetic activation of NDNF positive interneurons in anesthetized mice reduces spontaneous and evoked activity of excitatory neurons, suggesting an inhibitory effect, while chemogenetic suppression of NDNF positive interneurons reduces scaling of excitatory visual responses with pupil diameter, suggesting a disinhibitory effect^27^. Silencing LAMP5 positive interneurons was also shown to increase the spontaneous activity of excitatory neurons during locomotion, contributing to a lower visually evoked change in excitatory activity from pre- to post-onset^28^. While across the gamut of possible configurations of behavioral states, internal states, and sensory input either VIP or NDNF/LAMP5 positive interneuron activity may be more important to modulating a specific measure of excitatory activity, neither is likely to be wholly dispensable. Indeed, a screen across the spectrum of supragranular inhibitory interneurons suggests that behavioral state modulation exists as a transcriptomic spectrum, as opposed to being clearly delineated by discrete subtypes^16^.

Our finding that NDNF positive interneuron activity was highly correlated to running speed agrees with a recent study from the mouse auditory cortex, which found that NDNF positive interneurons were more correlated, time-locked, and responsive than VIP positive interneurons to running speed^29^. NDNF positive interneurons also received more pre-synaptic input from the motor cortex than VIP positive interneurons did^29^. While we similarly observed that changes in VIP positive interneuron activity lagged changes in running speed, we did not observe a significant lag in excitatory activity relative to running speed as was seen in the auditory cortex. This difference may relate to the effects of locomotion being largely repressive of auditory responses^40^. The layer specific differences we observe in NDNF positive interneurons add to a growing literature of depth dependent, rather than transcriptionally dependent, activity. NDNF/LAMP5 positive interneuron inputs, morphology, and transcriptomic subtype varies by depth. LAMP5 positive interneurons in layer 1 project horizontally while in layer 2/3 they are more spherical^31^. Within layer 1, NDNF/NPY double positive interneurons have a neurogliaform morphology with short dendrites and a wider axonal arbor, but NDNF positive NPY negative interneurons, termed canopy cells, have an even wider (but less tortuous) axonal arbor biased towards upper layer 1, and a higher density of boutons^21^. Functionally, putative neurogliaform cells were found to be more negatively modulated by behavioral state than putative canopy cells, whose activity was more closely aligned to VIP positive interneurons^16^. LAMP5 positive layer 2/3 neurons more uniformly increase activity in response to locomotion than LAMP5 positive layer 1 neurons^28^, which is similar to what we observed. Thus, our findings contribute insights to an evolving literature suggesting that VIP-to-NDNF state-dependent correlations vary transcriptionally, morphologically, and by cortical depth.

Across all three co-imaged pairs of populations, we observed overall positive correlations at the mean and pairwise level. The high pairwise correlation we observed between VIP positive interneurons and excitatory neurons corroborates previous reports^14,18^. The high VIP-to-NDNF correlation is consistent with their overall similarity in behavioral and visual drive, while the depth-dependent correlation may reflect separate subpopulations of NDNF positive interneurons, as discussed above. The increase in excitatory-to-VIP and decrease in excitatory-to-NDNF correlation during locomotion is intriguing and may reflect the prolonged activity of NDNF positive interneurons following locomotion onset. It should be noted that our interpopulation results contrast sharply with a study using voltage indicators to simultaneously image multiple neuronal subtypes in mouse visual cortex^41^. However, several findings from that study run counter to the broader literature. Using voltage imaging, they observed that SOM positive interneuron activity increased more than VIP positive interneuron activity during arousal and following air puff, with 34% of VIP positive interneurons suppressed by air puff. This contrasts sharply with other studies, even in the absence of visual stimuli or with recordings using in vivo patch clamp electrophysiology as opposed to calcium imaging, which are two potential methodological differences^7,9,13,15,17^. They reported low NDNF-to-NDNF pairwise correlation, whereas previous studies report the NDNF positive interneuron population to be highly synchronous^27^. They reported very low correlation between VIP positive interneurons and excitatory neurons, which have been previously observed to have very high correlation^14,18^. Finally, they reported virtually uncorrelated dynamics between VIP positive and NDNF positive interneurons, whereas we observe VIP-to-NDNF correlations to be higher than between any other pair of cell types. As the prevalence of voltage imaging increases, this divergence between calcium and voltage activity should be reconciled.

The increase in the amplitude of excitatory visual responses during locomotion is now well validated^1,15^. However, there is an increasingly nuanced view of how the motoric and autonomic driven components of locomotion contribute to increasing visual responses, and how locomotion alters different aspects of visual encoding. Locomotion does not seem to alter orientation selectivity or preference while increasing activity uniformly across contrasts; however, the gain appears relatively higher for cells preferring higher spatial frequencies^2,4,5^. As VIP and NDNF positive interneurons are suggested to preferentially impact low contrast perception, we had hypothesized that there would be a selective effect of VIP and NDNF activity in predicting response amplitude for low contrast stimuli, however our largest effect sizes were in fact for high contrast stimuli for both behavior and interneuron activity. The higher gain of locomotion for high spatial frequency preferring neurons is likely due to increased VIP activity, as has been demonstrated in a series of studies related to the role of VIP and SOM positive interneurons in spatial frequency and size tuning^42–45^. Similar to previous studies, we distinguish the effects of arousal and locomotion by comparing pupil dilations in the absence of locomotion separately from locomotion, with pupil dilations in the absence of locomotion generally being lower amplitude and alternatively termed vigilance or quiet wakefulness^3,7^. Previous studies have found that arousal suppresses spontaneous activity, increasing the signal-to-noise of visual responses, while locomotion increases spontaneous firing leading to a multiplicative gain for quiet wakefulness (response to preferred stimulus increases more than non-preferred) and additive for locomotion (both preferred and non-preferred response increases)^3,7^. Despite using very similar visual stimuli, we observed modest, stimulus-dependent increases during vigilance (most evident for natural movies) and clear multiplicative scaling during locomotion. Our results are more in line with a study analyzing the Allen Brain Observatory dataset, which did not find an effect of pupil diameter on decoding during stationary periods and that the primary effect of locomotion was on the response to the preferred stimulus^5^. Given the high correlation of running speed and pupil diameter, and the fact that pupil diameter is only an indirect readout of a broader physiological state of arousal, the differing results may reflect differences in criteria or recording method. Meta studies across multiple datasets are increasingly feasible with the increased acceptance of open science standards, and it is likely this ambiguity will be resolved in the near future.

In conclusion, we find that within layer 2/3 of the mouse visual cortex VIP positive interneurons are more correlated to pupil diameter than are NDNF positive interneurons, while both NDNF and VIP positive interneurons are correlated to locomotion. Locomotion increases excitatory-to-VIP correlation while decreasing excitatory-to-NDNF correlation. VIP and NDNF positive interneurons have distance dependent correlation to excitatory neurons and to one another, however VIP-to-NDNF correlation varies with cortical depth, as does the correlation of NDNF positive interneurons to behavioral state. Behavioral state and contrast are both powerful modulators of excitatory visual responses, however the activity of both VIP and NDNF positive interneurons preceding stimulus onset also contributes positively to the amplitude of excitatory visual responses beyond their correlation to behavioral state. Thus, while functionally similar, VIP and NDNF positive interneurons contribute distinct state-dependent signals.

## Author Contributions

This project was conceptualized by M.S., K.J., S.A.R., and G.S. All data collection was performed by K.J. Analysis was performed by K.J. with help and feedback from S.A.R., M.C., and G.S. Writing was done by K.J. with input from all authors. All authors read and approve the final manuscript.

## Acknowledgements

This work was supported by NIH grants R01MH126351 (M.S.), R01EY028219 (M.S.), 1K99EY035752 (S.A.R.), and F32EY032756 (K.J.). In addition, this work was supported by the Picower Institute Innovation Fund (M.S.) and the European Union’s Horizon research and innovation programme grant agreement No. 101206609, Marie Sklodowska-Curie Global Fellowship - AstroError (M.C.). The authors thank Taylor Johns and other members of the Sur lab for their help and support.

## Methods

### Experimental model and subject details

All procedures were approved by the Massachusetts Institute of Technology Animal Care and Use Committee (protocol 2308000562) and conformed to NIH guidelines. We used male and female mice. Mice were group housed with littermates after weaning (no more than 5 per cage) and single housed after surgery. Mice had a 12/12-hour reversed light/dark cycle, an average temperature of 21 °C, and an average humidity of 50%. We recorded during the dark cycle. Food and water were available ad libitum. We used the B6J.Cg-Viptm1(cre)Zjh/AreckJ (VIP-Cre, RRID:IMSR_JAX:031628), Ndnftm1.1(cre)Rudy/J (NDNF-Cre, RRID:IMSR_JAX:030757), and Viptm2.1(flpo)Zjh/J (VIP-Flp, RRID:IMSR_JAX:028578) lines or their crosses. All mice used were heterozygous for the transgenes. In the VIP-Cre cohort (VIP-Cre mice), we co-imaged VIP interneurons and putative excitatory neurons (157 recordings from 10 mice). In the NDNF-Cre cohort (NDNF-Cre mice), we co-imaged NDNF interneurons and putative excitatory neurons (114 recordings from 7 mice). In the VIP-Flp x NDNF-Cre cohort (VIP-Flp x NDNF-Cre mice), we co-imaged VIP and NDNF interneurons (17 recordings from 4 mice).

### Surgical preparation

We performed surgeries in adult mice. We anesthetized mice with isoflurane (2%) and gave pre-surgical slow-release buprenorphine (1 mg/kg) and meloxicam (5 mg/kg) subcutaneously. We placed the mouse on a Kopf stereotax and maintained body temperature with a heating pad (ATC2000, World Precision Instruments). We removed fur from the scalp with Nair, injected 0.1 mL of bupivacaine (0.5%) locally, and cleaned the scalp with 3 alternating applications of betadine and 70% ethanol. After confirming appropriate anesthesia, we removed the scalp with iris scissors to expose the skull over the frontal and interparietal bones. We leveled the skull, scraped it clean, and scored it with a scalpel. We outlined a 3 mm circle on the left hemisphere, 1 mm anterior and 3 mm lateral to the intersection of the lambda and midline sutures. We performed a circular craniotomy while frequently wetting the skull with ice-cold mammalian Ringer’s solution (50-980-245, Fisher Scientific), until we could lift the bone fragment under a drop of Ringer’s. We injected adeno-associated viruses (AAVs) diluted in Ringer’s solution (QSI 53311, Stoelting) through a pulled glass pipette with a beveled 50 µm opening, 0.2 to 0.4 mm below the dura, at 3 to 4 locations spaced more than 0.5 mm apart. Injection sites were 2.5 to 3.5 mm lateral and 0 to 1 mm anterior to the intersection of the lambda and midline sutures, which targets the monocular region of left primary visual cortex. We injected 150 nL per site at 50 nL/min.

In the VIP-Cre and NDNF-Cre cohorts, we injected AAV9-syn-jGCaMP7f-WPRE (104488, Addgene) and rAAV1-Flex-tdTomato (lot AV5328A2, UNC Vector Core). In the VIP-Flp x NDNF-Cre cohort, we injected AAV1-Flex-jRGECO1a (100853-AAV1) and AAV1-hEf1a-fDIO-rev-GCaMP6f (lot 150399, Virotek). We sealed the craniotomy with a stack of two 3 mm circular coverslips and one 5 mm circular coverslip (CS-3R and CS-5R, #1 thickness, Warner Instruments), glued together with optical adhesive (NOA 61, Norland), which compressed the cortex and improved imaging stability. We covered the edge of the 5 mm coverslip and the exposed skull with Metabond (C&B Metabond, Parkell) mixed with black ink pigment (Black Iron Oxide 18727, Schmincke) to limit light leak. We fixed a custom stainless-steel headplate (eMachine Shop) over the window with more Metabond. Mice recovered from anesthesia in a clean cage with heat, saline, and additional meloxicam as needed during a 3-day recovery period. Imaging began at least 2 weeks after window implantation.

### Behavioral recording and visual stimuli

We head-fixed mice on a running wheel (Bio-Serv) attached to a rotary encoder (LPD3806-600BM-G5-24C). A monitor (15 x 9 cm, 800 x 600 pixels) was placed perpendicular to the mouse and 14 cm in front of the eye contralateral to the imaged hemisphere. We displayed drifting gratings (1 second duration, 0.04 cycles per degree, 2 cycles per second, 8 directions in 45-degree steps) or natural movies (2 second duration) at multiple contrasts with a 4 second interstimulus interval^46^. Presentation order of each unique stimulus/contrast pairing was randomized across 10 blocks. A custom MATLAB script controlled stimulus timing, the air puff, and wheel acquisition; we generated visual stimuli with Psychtoolbox and logged wheel rotation through a microcontroller board (Arduino UNO). The initial cohorts of VIP-Cre and NDNF-Cre mice used varied contrasts, so for consistency when visual responses are reported only sessions from mice where 4%, 16%, and 64% contrast were used are reported. In some sessions, we delivered a mild air puff to the whisker pad and eye ipsilateral to the imaged hemisphere through a small tube 3 cm away (compressed air at 40 psi for 0.3 s; solenoid valve 003-0137-900, Parker; Arduino UNO controller). The puff ended 2 s before the onset of the 90-degree grating or movie 1, and we alternated blocks of puff and no-puff presentations of that stimulus, so that visual responses preceded by puff could be excluded from calculations of visual responsiveness. We recorded the contralateral face and pupil with a Thorlabs CMOS camera (CS165MU Zelux) and a 25 mm C Series fixed focal length lens (#59-871, Edmund Optics). A light-emitting diode array (LIU780A, Thorlabs; 780 nm) provided infrared illumination. A custom MATLAB script acquired video at 20 Hz. A National Instruments data acquisition device (BNC-2110 terminal block) generated 5-volt square-wave pulses that aligned the stimulus presentation, pupil imaging, and calcium imaging.

### Two-photon imaging

We imaged with a Prairie Ultima IV two-photon microscope using resonant-galvo scanning (Prairie View 5.4.64.500) and a 20x objective (XLPlan N 20x 1.00 NA, Olympus) at 2x digital zoom. Two-photon excitation came from a tunable laser (MaiTai eHP or Insight X3+, Spectra-Physics) at 900-920 nm in the VIP-Cre and NDNF-Cre cohorts and 965 nm in the VIP-Flp x NDNF-Cre cohort. Power at the objective ranged from 10 to 30 mW depending on depth and expression. The resonant-galvo scanner acquired frames at approximately 30 Hz (frame period 0.0328 s) at 512×512 pixels. For single-plane imaging, every 4 frames were averaged, giving an effective rate of 7.63 Hz. For dual-plane imaging with a piezo (17 of 114 NDNF-Cre recordings and 9 of 17 VIP-Flp x NDNF-Cre recordings), we imaged at a resolution of 512×256 with 4 frame averaging for an effective rate of 5 Hz. To account for offset in the plane of imaging relative to the cortical surface, depth was set relative to the cortical surface at the center of the field of view.

### Preprocessing of behavioral data

#### Pupil

We used DeepLabCut to extract 8 xy-coordinates around the pupil^47^. We trained a separate classifier for each mouse. We computed pupil diameter as the horizontal distance between two opposing markers scaled by the median ratio of vertical to horizontal diameter, and converted it to millimeters with a camera calibration of 60.8 pixels/mm. We replaced artifacts such as blinks (frames whose frame-to-frame change fell below the 5th percentile of negative changes) with modified Akima interpolation, resampled the trace to 20 Hz by linear interpolation, filled values of 2.5 mm or greater and missing values with the previous value, and smoothed with a 1 s moving median. In cases where frames were dropped during recording, we estimated frame timestamps with linear spacing. We defined pupil dilation events with four criteria. The smoothed diameter must have a forward 1 s moving mean at least 1.1 times the mean of the preceding 2 s. The velocity must be at least 10 µm/s. Events must last at least 1 s. Events must be separated by at least 2 s. A dilation ended (constriction) when the diameter fell to half of the peak amplitude above the preceding baseline while velocity was non-negative; overlapping dilations not separated by at least 2 seconds were merged.

#### Locomotion

We converted encoder counts to speed in cm/s with an empirically determined factor of 0.0158 cm per count over 0.02 s. We took the absolute value, interpolated linearly to 20 Hz, and smoothed with a causal moving median over the preceding 0.5 s. We defined locomotion events as periods in which smoothed speed stayed at or above 1 cm/s for at least 1 s, with at least 2 s between events.

### Behavioral state segmentation

We assigned every 20 Hz sample to one of three non-overlapping behavioral states:

- Quiescence: no pupil dilation event and no locomotion event.
- Vigilance: the sample lies inside a pupil dilation event and outside any locomotion event, excluding dilation periods that began while the mouse was locomoting.
- Locomotion: the sample lies inside a locomotion event.

Assigning state requires wheel data, so only recordings with a valid wheel trace contribute to state-based analyses. N therefore differs across analyses. Recordings without wheel data contribute to pupil-based analyses only. We assigned each visual trial to the state at the sample nearest the stimulus onset. For analyses of pupil dilation not accompanied by locomotion (**Figure 2D**), we used dilations with no locomotion sample within 1 s before or after onset.

### Calcium processing and spike inference

We detected ROIs with Suite2p (version 0.10.3; Cellpose detection with a cell diameter of 25 pixels)^48^. We corrected for neuropil contamination as F_corrected_ = F_raw_ − 0.7 × F_neuropil_. We then calculated dF/F = (F − F0)/F0, where F0 was the mean fluorescence in the second half of each interstimulus interval. We inferred spiking with Cascade applied to the dF/F traces, using the pretrained model whose training frame rate was nearest to the effective frame rate of the recording (5 or 7.5 Hz)^32^. Excitatory neurons used the global excitatory models, and VIP and NDNF interneurons used models trained on a VIP dataset (DS26-GCaMP6f-m-VIP-V1). Cascade returns the number of spikes in each imaging frame; we multiplied this by the frame rate to give an inferred firing rate in spikes per second. We then interpolated firing rates to the 20 Hz behavioral time base. Where a figure shows z-scored activity, we subtracted each neuron’s mean firing rate over the whole recording and divided by its standard deviation.

### Cell-type identification, inclusion and exclusion

We identified VIP and NDNF interneurons by their expression of the Cre- or Flp-dependent red fluorescent reporter (tdTomato in VIP-Cre and NDNF-Cre; jRGECO1a in VIP-Flp x NDNF-Cre). We selected these ROIs by manual curation in Suite2p (see Calcium processing). We refer to the red-negative, syn-GCaMP7f-expressing ROIs as putative excitatory neurons. Because the synapsin promoter drives expression in all neurons, this population can include unlabeled interneurons^31^. To limit contamination of the excitatory population by unlabeled interneurons, we kept excitatory neurons only from recordings with imaging-plane depth of at least 100 µm. Every panel that shows excitatory neurons or activity therefore uses planes deeper than 100 µm. In the VIP-Flp x NDNF-Cre recordings, double-labeled neurons were rejected^21^. We required at least 4 excitatory neurons and at least 1 interneuron per recording for correlations between populations. For state-resolved analyses, we required that each of the three states occupy at least 5% of the recording. Cross-covariance peaks at the first or last lag of the window were excluded from the lag, peak, and asymmetry estimates.

We called a neuron visually responsive when a permutation test of its response (mean activity during the stimulus minus mean activity in the second half of the preceding interstimulus interval) to its preferred grating or movie at the highest contrast gave p < 0.05. The test compared the observed mean response with a null distribution built from 10,000 random circular shifts of the stimulus times. As this analysis was only run once per neuron (on its preferred stimulus at 64% contrast), we did not correct the responsiveness test for multiple comparisons across neurons. Responsiveness to pupil dilation and locomotion onset used the same permutation test (10,000 shifts) on activity in the 5 s after versus the 5 s before event onset, with p < 0.05. This was only performed in sessions with more than two pupil dilation or locomotion onset events, and only these sessions were used in calculating the fraction responsive to visual stimuli, pupil dilation, and locomotion onset. We excluded trials of the puff stimulus that were delivered during puff blocks from the state-resolved visual responses. Only sessions using 4%, 16%, and 64% contrast were used for visual response analyses for consistency between cohorts.

### Visual response measures

For each trial we calculated the response amplitude as the mean firing rate during the stimulus minus the mean firing rate in the second half of the preceding interstimulus interval. We averaged trials of the same stimulus and contrast. A neuron’s preferred direction (or preferred movie) was the stimulus with the largest mean amplitude at the highest contrast. For gratings, we aligned responses to the preferred direction by circularly shifting the 8 directions and called the direction at 0 degrees the preferred direction and the two directions at plus and minus 90 degrees the orthogonal directions. In preferred-versus-orthogonal analyses, orthogonal response is the mean of those two directions. Normalized responses were divided by the neuron’s response to its preferred stimulus at the highest contrast across all states. **Figure 4E** shows responses to absolute screen directions rather than preferred-aligned directions.

For frames in the quiescent state, we averaged the firing rate of all co-imaged interneurons at every 20 Hz time point. We then computed the 25th, 50th, and 75th percentiles of the nonzero values and assigned each time point to a quartile (q1 to q4). We assigned each visual trial to the bin at the sample nearest stimulus onset. **Figure 5A-H** shows trials only during quiescence to best isolate the unique contribution of preceding interneuron activity to excitatory response amplitude in the absence of elevated behavioral state.

### Measures of neuronal activity, behavior, and coupling

#### Correlation to pupil and to running speed

For each neuron we calculated the Pearson correlation between its firing rate and the smoothed pupil diameter or running speed, then subtracted the mean correlation obtained from 1,000 random circular shifts of one trace. Partial correlations used MATLAB partialcorr, controlling for the other behavioral variable.

#### Cross covariance and lag

We calculated cross covariance with xcov and the ‘normalized’ option, which scales the value at zero lag to the Pearson correlation, at 20 Hz. For a neuron and a behavioral trace, the lag at the maximum is the lag at which the neuron best matches the behavior; a positive lag means the neuron’s activity follows the behavioral signal (xcov(neuron, behavior)). Peaks at the first or last lag were excluded. We calculated population cross covariance from the mean activity of each population in one recording. In the state-resolved analyses, we calculated covariance within each contiguous bout of a state that lasted at least 2.05 s and averaged the covariance across bouts. For the pairs of populations, the first signal is excitatory neurons in VIP-Cre and NDNF-Cre and VIP interneurons in VIP-Flp x NDNF-Cre; a negative lag means that the first population changed before the second.

#### Correlation to the other population

For each neuron we averaged its activity-adjusted Pearson correlations (1,000 circular shifts) with all co-imaged neurons of the other population, overall and within each state.

#### Distance dependence

We converted ROI centroids to micrometers and calculated Euclidean distances between neurons of the two populations in the same recording. Distances were binned in 50 µm bins from 0 to 500 µm.

#### Event-aligned responses

We aligned z-scored activity to the onset of pupil dilations (without locomotion), locomotion onset, or air puffs and averaged across events. The response amplitude is the mean in the 5 s after onset (0 to 5 s) minus the mean in the 2 s before (−2 to 0 s).

#### Deciles

We calculated the 10th to 90th percentiles of pupil diameter and running speed for each recording, took the median across a mouse’s recordings, and used these values as bin edges for that mouse. Neuronal activity was then averaged in each of the 10 bins.

### Statistical analysis

#### General

We analyzed data with linear mixed-effects models (MATLAB fitlme, fit by maximum likelihood). Random effects always included the subject ID. For neuron-level data they also included the recording ID, and neuron ID if repeated measures were taken for each neuron (e.g., mean activity by behavioral state). We tested each fixed effect with F-tests from anova(model, ‘DFMethod’, ‘satterthwaite’), which uses Satterthwaite degrees of freedom (non-integer, reported as F(df1, df2)). For significant fixed effects, we then tested single coefficients with the t stat that fitlme reports; their degrees of freedom are residual degrees of freedom (number of observations minus number of fixed effects). In models with an interaction, a single coefficient is the simple effect relative to the reference level of the other terms. Reference levels were quiescence for state, excitatory neurons for cell type, and the first level of each categorical variable otherwise. For the comparison of VIP with NDNF we refitted the model with VIP as the reference level.

#### Post hoc tests

For the effect of behavioral state within each contrast, we fitted a separate model to the data from each contrast and report the coefficient t stat for vigilance and locomotion versus quiescence. For the effect of direction (**Figure 4E**) and the effect of interneuron quartile at each contrast (**Figure 5**), we likewise fitted separate models. We did not correct these post hoc tests, the coefficient tests of cell type or state, or the signed-rank tests for multiple comparisons.

#### Signed-rank tests

We tested lags and cross-covariance peaks with two-sided Wilcoxon signed-rank tests using the normal approximation (signrank with method ‘approximate’); we report Z and the number of observations (neurons or recordings). The unit of observation is stated for each result.

#### Permutation tests of distance dependence (Figure 3G)

For each cohort, we pooled the pairs of neurons in all recordings and binned the pairwise correlations by distance. We shuffled the neuron identities of one population within each recording (10,000 shuffles) to generate a null distribution of mean correlation for each bin. The p-value of a bin was twice the smaller of the proportions of null means above and below the observed mean, multiplied by the number of bins (10) as a Bonferroni correction, and not capped at 1. With 10,000 shuffles and 10 bins, the smallest p-value that the test can resolve is 2 x 10 / 10,000 = 2e-3 (computed).

#### Single-trial models and cross-validation (Figure 5I-L; Figure S4)

For each visually responsive excitatory neuron, we used its trials at its preferred stimulus (all three contrasts) and calculated four trial-level variables: response amplitude (stimulus minus preceding interstimulus activity), mean pupil diameter (pupil), mean running speed (speed), and mean firing rate of the co-imaged interneurons (inter: VIP in VIP-Cre, NDNF in NDNF-Cre). We calculated the latter three variables over the 0.5 s before stimulus onset. We z-scored the response amplitude and the three predictors across trials, and z-scored the natural logarithm of contrast. We kept trials with all variables present and mice that had exactly the contrasts 4%, 16%, and 64%. The full model was trial_amp ~ contrast*(pupil + speed + inter) + (1|subject) + (1|session) + (1|neuron_id). The reduced models were: contrast only; contrast with one predictor and its interaction with contrast; and the full model with one term removed (contrast with its interactions; pupil; wheel; or interneuron activity). We estimated explained variance by 100 repartitions of 3-fold cross-validation over trials (cvpartition ‘KFold’). In each fold, we fitted the model to the training trials and predicted the held-out trials. Held-out R-squared is 1 minus the sum of squared prediction errors divided by the total sum of squares of the held-out trials around their mean. The marginal contribution of a predictor is the R-squared of the model with the predictor (and its interaction with contrast) minus the R-squared of the contrast-only model. The unique contribution of a predictor is the R-squared of the full model minus the R-squared of the full model with that predictor (and its interaction with contrast) removed. The distributions over the 100 repartitions are shown. We tested significance on the full dataset (not on the cross-validated values) with likelihood-ratio tests with MATLAB’s compare function and report chi-square, degrees of freedom, and p. We estimated p-values that the software printed as 0 as p < 1e-300, estimated given the precision of the float type. Single-predictor models were tested against the contrast-only model (marginal) and reduced models against the full model (unique).

