## Supplemental Figures 1-4 for "Correlated but distinct: VIP and NDNF interneurons implement dissociable inhibitory motifs for state-dependent gain control in visual cortex"

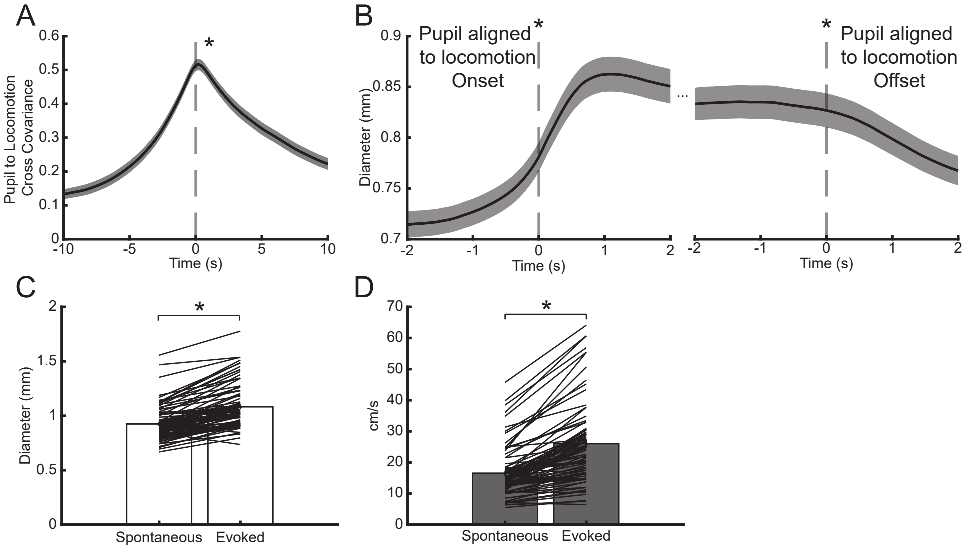


**Supplemental Figure 1: Pupil diameter and locomotion are correlated but offset. A.** Cross covariance of pupil diameter and running speed. Pupil diameter and running speed significantly covary (z(145) = 10.426, p = 1.88e-25, signed rank test versus zero). Changes in running speed significantly preceded changes in pupil diameter by 0.33 seconds (z(145) = 4.509, p = 6.52e-6, signed rank test vs. lag of zero seconds). **B.** Pupil diameter aligned to locomotion onset (left) and offset (right). Pupil diameter significantly increases at locomotion onset (z(145) = -10.444, p = 1.56e-25, signed rank test) and decreases at locomotion offset (z(145) = 9.319, p = 1.17e-20, signed rank test). **C.** Pupil dilations evoked by puffs are significantly larger than spontaneous pupil dilations (z(86) = -7.87, p = 3.45e-15, signed rank test). **D.** Bouts of locomotion evoked by puff are significantly faster than spontaneous bouts of locomotion (z(85) = -7.938, p = 2.06e-15, signed rank test). *denotes significance.

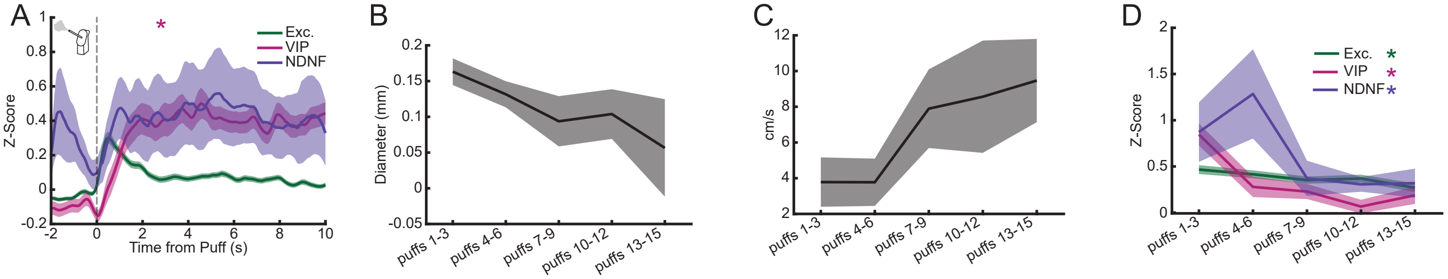


**Supplemental Figure 2: Behavioral and neuronal responses to air puffs attenuate over time. A.** Averaged Z-scored neuronal activity aligned to air puff. Response amplitude varies by cell type ($F(2,1113.2) = 6.060$, $p = 2.41e\text{-}3$, ANOVA), with VIP ($t(1119) = 3.474$, $p = 5.32e\text{-}4$) responding with a higher amplitude than do excitatory neurons. **B.** The size of the pupil following air puff decreases over the length of the first session with air puff ($F(1,32) = 9.769$, $p = 3.76e\text{-}3$, ANOVA). **C.** Running speed following air puff increases over the length of the first session with air puff ($F(1,32) = 13.399$, $p = 8.99e\text{-}4$, ANOVA). **D.** Averaged Z-scored responses of excitatory neurons, VIP interneurons, and NDNF interneurons to air puff decrease over the length of a recording (Exc.: $F(1,986.65) = 17.991$, $p = 2.43e\text{-}5$. VIP: $F(1,114.68) = 40.588$, $p = 4.05e\text{-}9$. NDNF: $F(1,52) = 8.9446$, $p = 4.25e\text{-}3$, ANOVA). *denotes significance.

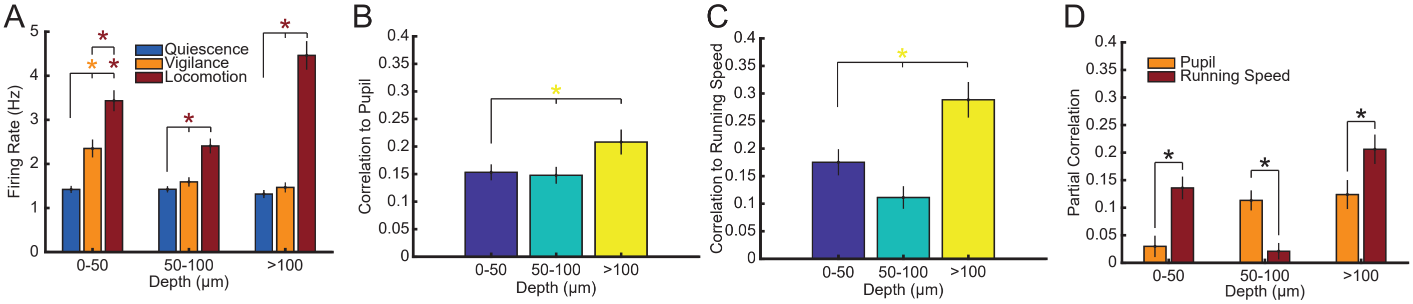


**Supplemental Figure 3: The response of NDNF interneurons to behavioral state varies by depth. A.** Average NDNF interneuron firing rate during each behavioral state varies by depth from the cortical surface (F(4,466) = 17.343, p = 2.83e-13, ANOVA state-depth category interaction). For 0-50 μm, NDNF firing rates during vigilance and locomotion are both higher than during quiescence (Vigilance: p = 4.24e-5. Locomotion: p = 7.69e-17. Vigilance vs. locomotion: p = 2.15e-6). However, for both 50-100 and >100 μm firing rate only increases significantly during locomotion (50-100 μm: quiescence vs. locomotion, p = 6.22e-10; vigilance vs. locomotion, p = 2.12e-7. >100 μm: quiescence vs. locomotion, p = 4.03e-22; vigilance vs. locomotion, p = 1.48e-20). **B.** NDNF interneurons' correlation to the pupil diameter varies by depth from the cortical surface (F(2,100.17) = 3.572, p = 0.0317, ANOVA). NDNF interneurons > 100 μm in depth have significantly higher pupil correlation than shallower NDNF interneurons (Vs. 0-50 μm: p = 0.0138. Vs. 50-100 μm: p = 0.0359). **C.** NDNF interneurons' correlation to running speed varies by depth from the cortical surface (F(2,19.022) = 7.861, p = 3.25e-3, ANOVA). NDNF interneurons > 100 μm in depth have significantly higher running speed correlation than shallower NDNF interneurons (Vs. 0-50 μm: p = 9.13e-4. Vs. 50-100 μm: p = 2.86e-3). **D.** NDNF interneurons' partial correlation to pupil diameter and running speed varies by depth from the cortical surface. NDNF interneurons from 0-50 μm have higher partial correlation to running speed than to pupil diameter (F(1,146.27) = 14.581, p = 1.98e-4, ANOVA). NDNF interneurons from 50-100 μm surprisingly flip this relationship (F(1,176) = 15.477, p = 1.2e-4, ANOVA), while NDNF neurons > 100 μm from the surface flip back to have higher correlation to running speed (F(1,136.68) = 5.185, p = 0.0243, ANOVA). *denotes significance.

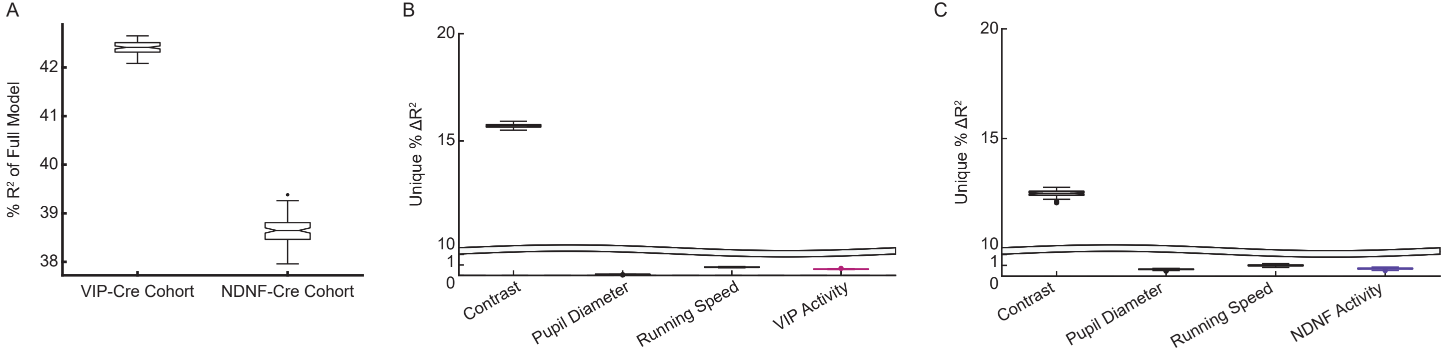


**Supplemental Figure 4: R-squared from full model with all predictors. A.** Mixed-effect linear models were fit to the amplitude of excitatory visual responses with contrast as a predictor, and the pupil diameter, running speed, and mean VIP or NDNF interneuron activity preceding stimulus onset (and their interaction with contrast) as predictors. R-squared of the full model across 100 rounds of three-fold cross validation in the VIP-Cre and NDNF-Cre cohorts. **B.** The unique variance explained was computed across 100 rounds of three-fold cross-validation by comparing the full model with reduced models that excluded, in turn, contrast, pupil diameter, running speed, or VIP activity preceding stimulus onset. The significance of the unique explained variance of each predictor compared to the full model was computed using the full dataset. Contrast, pupil diameter, running speed, and mean VIP activity preceding stimulus onset all have significant unique contributions to the explained variance (Contrast: $p < 1e\text{-}300$, likelihood ratio test). **C.** Same as B, but using mean NDNF activity preceding stimulus onset. Contrast, pupil diameter, running speed, and mean NDNF activity preceding stimulus onset all have significant unique contributions to the explained variance (contrast: $p < 1e\text{-}300$, likelihood ratio test). *denotes significance.
